# Bacterial glycoengineering as a biosynthetic route to customized glycomolecules

**DOI:** 10.1101/118224

**Authors:** Laura E. Yates, Dominic C. Mills, Matthew P. DeLisa

## Abstract

Bacteria have garnered increased interest in recent years as a platform for the biosynthesis of a variety of glycomolecules such as soluble oligosaccharides, surface-exposed carbohydrates and glycoproteins. The ability to flexibly engineer commonly used laboratory species such as *Escherichia coli* to efficiently synthesize non-native sugar structures by recombinant expression of enzymes from various carbohydrate biosynthesis pathways has allowed for the facile generation of important products such as conjugate vaccines, glycosylated outer membrane vesicles, and a variety of other research reagents for studying and understanding the role of glycans in living systems. This chapter highlights some of the key discoveries and technologies for equipping bacteria with the requisite biosynthetic machinery to generate such products. As the bacterial glyco-toolbox continues to grow, these technologies are expected to expand the range of glycomolecules produced recombinantly in bacterial systems, thereby opening up this platform to an even larger number of applications.

## Bacteria as a platform for polysaccharide and glycoconjugate production

In recent years, there has been growing interest in developing bacterial species as hosts for glycoengineering applications involving the biosynthesis of structurally diverse polysaccharides, which can be produced as free glycans or as conjugates to lipids or proteins. The most obvious advantage of this approach is the vastly simpler and cheaper culturing conditions required for maintenance of bacterial cells when compared to a eukaryotic cell culture. However, bacteria are in fact highly proficient producers of carbohydrates, with more than 140 unique monosaccharide base types identified in bacterial species, in contrast to the 14 base types produced by mammalian species (Herget et al. 2008). Many of these bacterial monosaccharides are then assembled into an even more diverse array of polysaccharides, often as part of surface structures such as capsular polysaccharide (CPS) and the O-antigen component of lipopolysaccharide (LPS), which are often important virulence factors in pathogenic species. In *Escherichia coli* alone, 187 unique O-antigen structures and 80 CPS structures have been identified to date (Stenutz et al. 2006; Senchenkova et al. 2016; Whitfield 2006). Other bacterial polysaccharides have important structural functions (e.g., peptidoglycan), or play a role in adaptation to environmental conditions by mechanisms such as osmoregulation (e.g., enterobacterial common antigen, ECA) (Kuhn et al. 1988).

The pathways responsible for production of mono- and polysaccharides are frequently well-defined in bacteria, especially in commonly used host species such as *E. coli* (Aoki-Kinoshita and Kanehisa 2015). Furthermore, with the exception of the ubiquitous structural polysaccharide peptidoglycan, bacterial polysaccharides are typically non-essential for viability, meaning biosynthesis pathways are amenable to genetic manipulation and deletion. For example, metabolic engineering studies have identified routes to enhance the availability of relevant nucleotide-activated sugars, leading to improved polysaccharide yields (Ruffing and Chen 2006). As a result of these and other related efforts, bacteria represent a tractable, well-defined platform for engineering the biosynthesis of polysaccharides.

While the ability of bacteria to produce polysaccharides and glycolipids is established, it was long believed that bacteria were incapable of modifying proteins with carbohydrate moieties, a process known as glycosylation. However, this paradigm was overturned in the 1970s with the identification of glycosylated surface layer (S-layer) proteins in *Halobacterium salinarum, Clostridium thermosaccharolyticum* and *Clostridium thermohydrosulfuricum* (Sleytr 1975; Sleytr and Thorne 1976). Although examples of bacterial protein glycosylation remain relatively uncommon, in the past 15 years a diverse array of systems have been discovered and characterized, including examples of sequential and *en bloc* transfer of both *N*-linked and *O*-linked glycans (Szymanski et al. 1999; Castric 1995; Grass et al. 2003; Thibault et al. 2001).

From an engineering perspective, perhaps the most significant advance came in 2002 with the functional transfer of a complete protein *N*-glycosylation system from the gastrointestinal pathogen *Campylobacter jejuni* into a laboratory strain of *E. coli*, which is naturally incapable of protein glycosylation (Wacker et al. 2002). The versatility of this system was further enhanced by a series of experiments demonstrating the modularity of the bacterial glycosylation machinery, which was found to tolerate a number of different glycan structures and protein substrates (Feldman et al. 2005; Kowarik et al. 2006; Wacker et al. 2006). Importantly, the newfound ability to generate glycoproteins in a genetically tractable host organism like *E. coli* provided a unique opportunity to both understand and exploit the glycosylation process in ways that were not previously possible with eukaryotic systems. This is because even though the pathways involved in the production of protein-linked polysaccharides in eukaryotic cells are well understood, the essential nature of many of these mechanisms limits the potential for manipulation.

### Polysaccharide production in bacteria

Enzymatic synthesis of polysaccharides utilizes nucleotide-activated sugars as glycosyl donors, to supply the necessary energy for the reaction. In bacteria, these nucleotide sugars are typically only present in the cytoplasm where they are synthesized. Consequently, all initial polysaccharide biosynthesis in bacteria also takes place within the cytoplasm. The majority of polysaccharides are synthesized by one of three pathways: the Wzy-dependent pathway, the ATP-binding cassette (ABC) transporter-dependent pathway, and the synthase-dependent pathway (Fig. 1), although shorter oligosaccharides may be formed by the direct action of glycosyltransferases on a substrate such as lipid A in the case of the LPS core or lipooligosaccharides (LOS) (Kalynych et al. 2014). The Wzy-dependent pathway involves the sequential action of glycosyltransferases on a lipid anchor, undecaprenyl diphosphate (Und-PP), on the inner leaflet of the cytoplasmic membrane, followed by translocation of a completed subunit across the membrane by the flippase Wzx. The subunits then undergo polymerization by the polymerase Wzy. The number of repeat units is modulated somewhat by Wzz, the chain-length regulator, although the resulting polymers are not strictly uniform in length. Completed polysaccharides are then removed from Und-PP and transferred to a target location, which differs depending on the species in question and the type of polysaccharide produced (Raetz and Whitfield 2002). Common examples of polysaccharides produced by this mechanism include the majority of O-antigen polysaccharides, and a significant proportion of capsules, as well as specific examples such as ECA, a surface polysaccharide common to most *Enterobacteriacae*, but limited to this family (Kuhn et al. 1988). In contrast, the ABC transporter-dependent pathway involves the assembly of the entire polysaccharide on a lipid anchor at the inner face of the cytoplasmic membrane, before the chain is capped to indicate completion, and the entire structure is transported across the membrane by the ABC-transporter complex (Cuthbertson et al. 2010). As with Wzy-dependent systems, however, the polysaccharide is then removed from the lipid anchor and transferred to a permanent point of attachment. Polysaccharides assembled by this method typically form O-antigen polysaccharides or capsules. Synthase-dependent polysaccharide assembly is unique in that it can occur in the presence or absence of a lipid anchor. A transmembrane glycosyltransferase simultaneously catalyzes formation of the polymer and translocation across the membrane (Whitney and Howell 2013). Polysaccharides produced by this mechanism may be attached to the exterior of the cell, but are more frequently released into the extracellular environment to form non-covalently associated exopolysaccharides such as hyaluronic acid (HA), alginate or cellulose.

**Figure 1.**
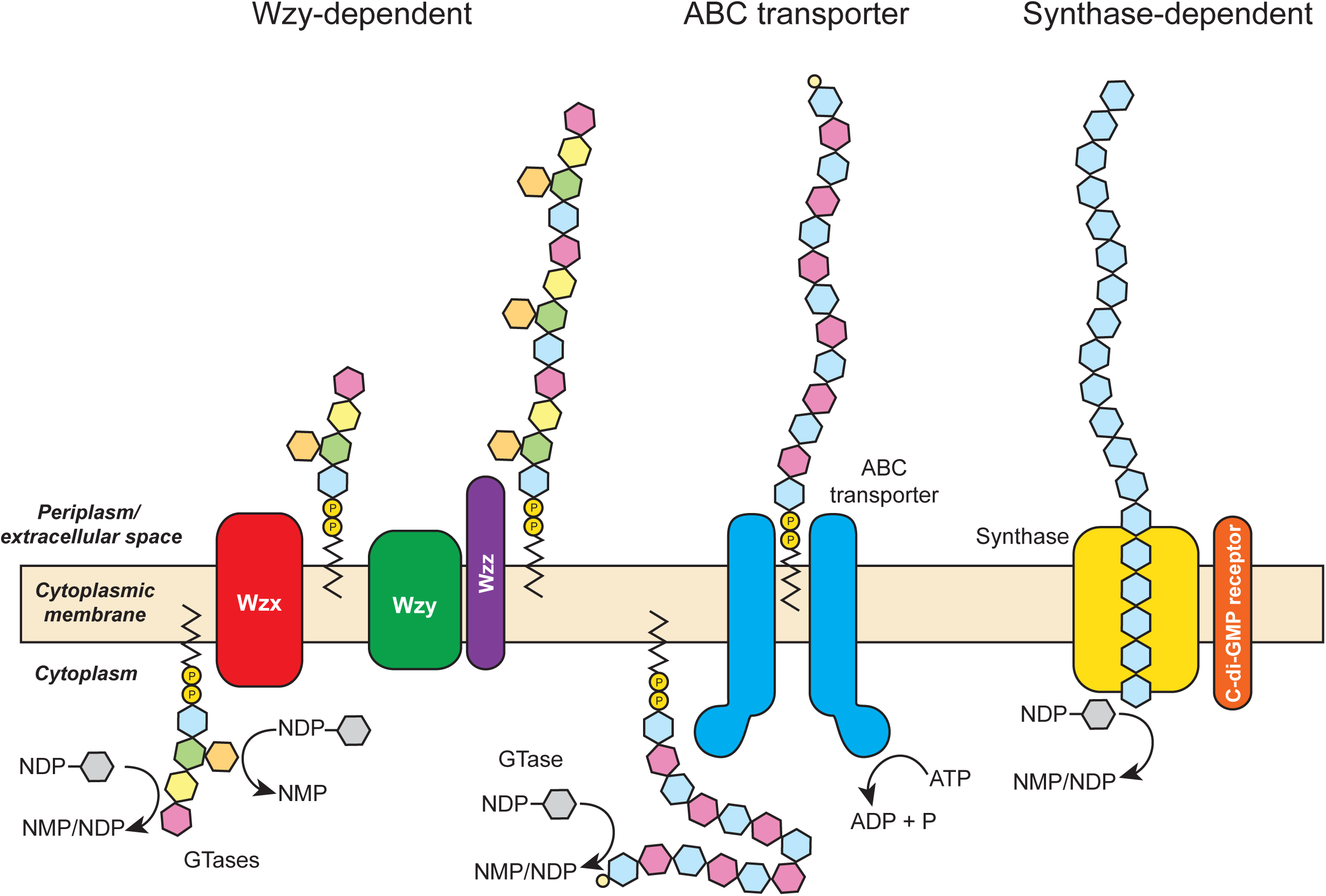
Biosynthesis of bacterial polysaccharides. The majority of bacterial polysaccharides are assembled by one of three mechanisms, the Wzy-dependent, the ABC transporter-dependent or the synthase-dependent pathway. The key protein components for each mechanism are indicated on the diagram, and are located in the inner membrane of Gramnegative organisms or the membrane of Gram-positive organisms. Polysaccharides are synthesized from nucleotide diphosphate (NDP) sugars. For the Wzy-dependent pathway, multiple glycosyltransferases (GTases) in the cytoplasm synthesize oligosaccharides on Und-PP. Oligosaccharides typically contain diverse monosaccharides and may be branched; consequently this assembly mechanism is responsible for the production of most high-complexity sugars. The completed oligosaccharide repeat unit is transported across the relevant membrane by the translocase or flippase enzyme Wzx. Multiple repeat units are then linked together by the polymerase enzyme Wzy to form a repeating heteropolymer. The final length of the polymer may be controlled by the chain length regulator Wzz. In the ABC transporter-dependent pathway, a homopolymer or simple heteropolymer is assembled on Und-PP on the cytoplasmic face of the membrane, often by just a single GTase. The completed polysaccharide is capped with a moiety such as a phosphate group, and transported through the membrane by the ATP-binding cassette (ABC) transporter. For synthase-dependent biosynthesis, the polysaccharide is simultaneously polymerized and transported across the membrane. In the absence of a membrane anchor, a receptor protein for a signaling molecule such as bis-(3′-5′)-cyclic dimeric guanosine monophosphate (c-di-GMP) may play a role in initiation of polysaccharide assembly. In Gram-negative organisms, polysaccharides are frequently transported across the outer membrane by an additional export system to enable surface display.

### Bioengineering of secreted oligosaccharides in bacteria

Small, soluble oligosaccharides play many important roles in biological systems, and as such have a multitude of potential uses in research, medicine and industry. However, owing to the extremely high heterogeneity of such structures, together with low yield and complex purification when isolating from natural sources, engineered production has been the focus of much research. Chemical synthesis is complex and costly, and the resulting oligosaccharides are subject to the same issues regarding heterogeneity, limiting their usefulness without significant downstream purification. Chemo- and in vitro-enzymatic methods have also been widely explored, and have shown great improvements with respect to yield and structural homogeneity, but isolation of the required enzymes is a demanding process, and the necessary nucleotide-activated sugars are extremely expensive to supply for such large-scale synthesis; consequently production beyond the milligram scale, especially for larger tri- and tetrasaccharides remains unfeasible by this method.

The development of a metabolically-engineered *E. coli* strain that could produce human milk oligosaccharides in a fermentation process represented a significant advance within the field (Priem et al. 2002). The engineered strain utilizes glycerol as an affordable carbon source, relying on native metabolic pathways within the bacterium to produce a continuous supply of the required nucleotide sugars. The approach also relies on the presence of a soluble acceptor sugar in the cytoplasm as an assembly platform. In this case lactose, which can be imported from the growth medium, was used. However, methods for the in situ synthesis of acceptor sugars have also been developed (Samain et al. 1997). Such engineered strains have been shown to produce quantities of up to 34 g/L of secreted oligosaccharide, and the scalable nature of production means the manufacture of kilogram quantities of sugar are entirely feasible (Drouillard et al. 2010). This approach has since been used for the production of more than 25 different oligosaccharides ranging from disaccharides to pentasaccharides, including structures that are known to have immunomodulatory effects or to be associated with cancer in humans (Ruffing and Chen 2006).

### Bioengineering of exopolysaccharides in bacteria

Many exopolysaccharides produced by bacteria have significant commercial value (Schmid et al. 2015), the most widely studied of which are listed in Table 1. Some of these polymers are naturally-occurring in bacteria, while others have been engineered via heterologous gene expression, particularly in cases where the original source or isolation method was undesirable. One example is HA, an extremely hydrophilic polymer of alternating β-D-glucuronic acid and β-D-*N*-acetyl-glucosamine residues that is a desirable material in medicine and cosmetics owing to its high water retention capacity and lack of toxicity. Initially, this polysaccharide was purified from rooster combs, although the majority of production is now achieved via microbial fermentation (Liu et al. 2011). Native bacterial production of HA was first achieved from *Streptococcus zooepidemicus* (Thonard et al. 1964), but due to co-production of the streptolysin exotoxin, recombinant production remained a priority. Indeed, recombinant HA was eventually achieved using the host organism *Bacillus subtilis* (Widner et al. 2005), and subsequently *E. coli* (Yu and Stephanopoulos 2008). Such approaches achieve yields of ~10 g/L, which is thought to be near the production limit owing to the effect of the exopolysaccharide on the viscosity of the growth medium (Yu and Stephanopoulos 2008). Key advances have come instead in the area of polymer length regulation, allowing for better control of physiochemical properties, and achieved largely through metabolic engineering and tighter control of the availability of the precursor nucleotide sugars (Jia et al. 2013).

**Table 1.** Extensively studied bacterial exopolysaccharides: composition, sources and uses.

| EPS | Components | Organism | Main applications* |
| --- | --- | --- | --- |
| Cellulose | Glucose | *Gluconacetobacter xylinus* | Foods (indigestible fiber) |
|  |  |  | Wound healing<br>Engineered blood vessels<br>Audio speaker diaphragms |
| Xanthan | Glucose<br>Mannose<br>Glucuronic acid<br>Acetate<br>Pyruvate | *Xanthomonas campestris* | Foods<br>Petroleum industry<br>Pharmaceuticals<br>Cosmetics and personal care products<br>Agriculture |
| Alginate | Guluronic acid<br>Mannuronic acid<br>Acetate | *Pseudomonas aeruginosa*,<br>*Azotobacter vinelandii* | Surgical dressings<br>Wound management<br>Controlled drug release |
| Gellan | Glucose<br>Rhamnose<br>Glucuronic acid<br>Acetate<br>Glycerate | *Sphingomonas paucimobilis* | Foods<br>Pet food<br>Pharmaceuticals<br>Agar substitute |
| Dextran | Glucose | *Leuconostoc mesenteroides* | Foods<br>Blood volume expander<br>Chromatographic media |
| Curdlan | Glucose | *Agrobacterium tumefaciens*, | Foods<br>Pharmaceuticals |
|  |  | *Alcaligenes faecalis* | Heavy metal removal<br>Concrete additive |
| Hyaluronic acid | Glucuronic acid<br>N-acetylglucosamine | *Streptococcus zooepidemicus*,<br>*Bacillus subtilis* | Medicine<br>Solid culture media |
| Succinoglycan | Glucose<br>Galactose<br>Acetate<br>Pyruvate<br>Succinate | *Sinorhizobium meliloti* | Food<br>Oil recovery |
| Levan | Fructose | *Bacillus subtilis*,<br>*Zymomonas mobilis* | Food (prebiotic)<br>Medicines<br>Cosmetics |

\*summarized from (Schmid et al. 2015)

In other cases, such as the commercially valuable xanthan, metabolic engineering has enabled yields of up to 50 g/L, also thought likely to be the highest level feasible for bioreactor processing (Seviour et al. 2011). Further increases will rely on additional engineering strategies to alter the molecular structure of the polysaccharide and reduce the resulting viscosity via modifications such as limiting polymer length or altering the degree of acylation or pyruvylation of a compound (Schmid et al. 2015). Bacterial production also offers unprecedented levels of purity when compared to extraction methods from other sources - for example, cellulose free from the common plant contaminants ligin and hemicellulose (Klemm et al. 2005). Furthermore, with the growing understanding of the pathways behind bacterial synthesis of such exopolysaccharides and recent advances in bioinformatics and systems biology, it may soon be possible to engineer bacteria to produce entirely novel polysaccharides with useful chemical properties. Indeed, a metabolic engineering approach was recently used to synthesize a variant form of cellulose containing a proportion of GlcNAc monomers in addition to the usual glucose. This modification resulted in the production of a biopolymer that is far more readily biodegradable than the standard form (Yadav et al. 2010).

### Bioengineering of intracellular and cell-associated polysaccharides in bacteria

The most widely manipulated cellular polysaccharide biosynthesis system is probably the LPS pathway (Fig. 2), in part due to the significance of this polysaccharide in pathogenesis, but also owing to the conserved mechanistic nature of the pathway combined with the highly variable glycan structures produced.

**Figure 2.**
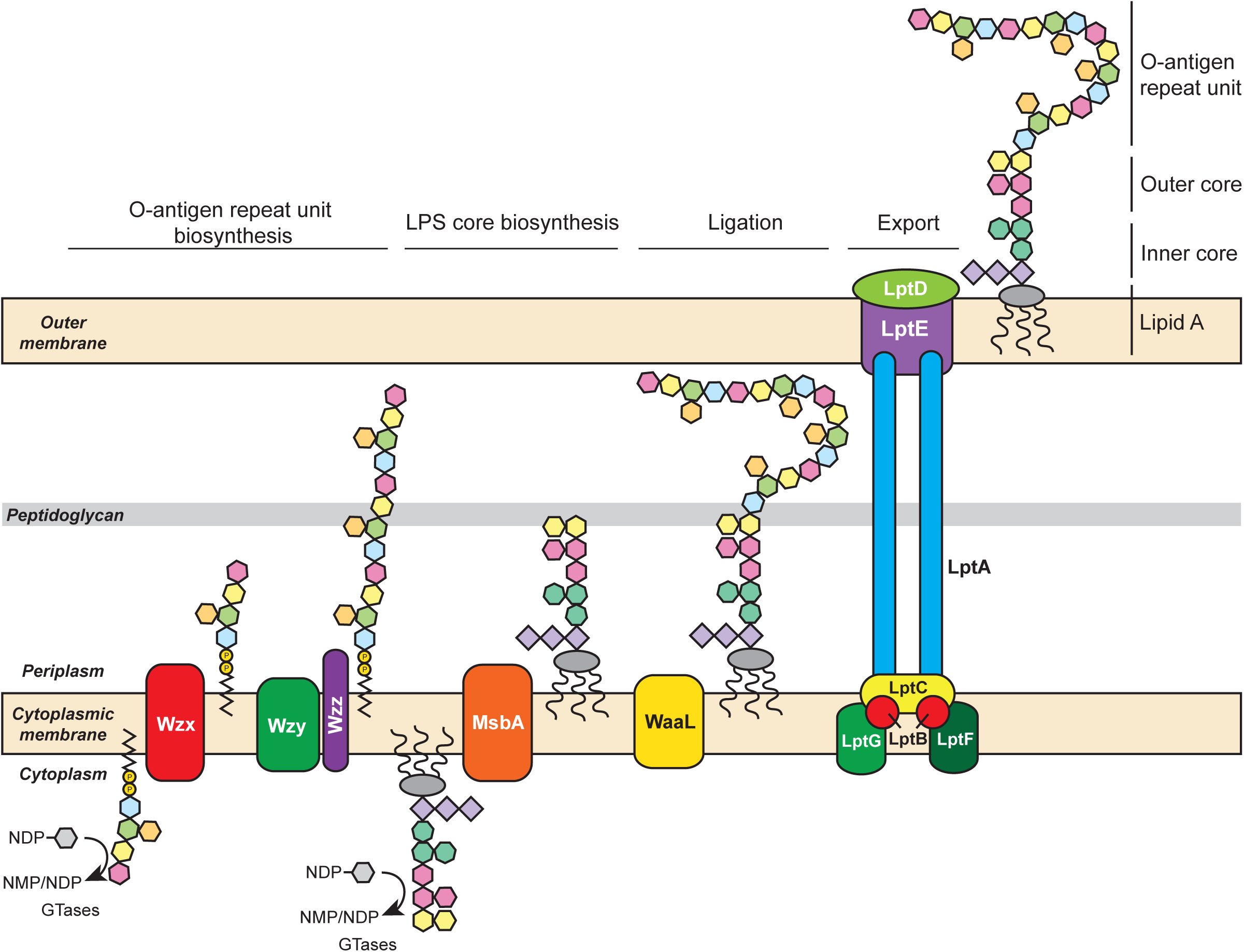
LPS biosynthesis. Multiple glycosyltransferases (GTases) transfer NDP-sugars to the nascent oligosaccharide to form an O-antigen repeat unit on Und-PP in the cytoplasm of the cell. The completed oligosaccharide is transported across the inner membrane by the flippase, Wzx, and multiple repeat units are joined together by the polymerase enzyme Wzy to form the completed O-antigen portion of the LPS. The structure shown is representative and does not indicate a specific serotype. Simultaneously, the LPS core, comprising lipid A, and the inner and outer core sugars is assembled in the cytoplasm. The inner core consists of two or three Kdo monosaccharides (diamond shapes), which are added during synthesis of the lipid A molecule, and three heptose sugars (heptagon shapes) which are added by the sequential action of three GTases. The outer core shown is an R1 structure, consisting of three glucose and two galactose residues (hexagons), and is assembled by the sequential action of a further five GTases. The completed LPS core is transported across the inner membrane by the ABC transporter MsbA. The O-antigen repeat unit is removed from the Und-PP membrane anchor and attached to the first galactose on the R1 outer core by the ligase enzyme WaaL. The entire LPS structure is then extracted from the inner membrane and transported across the periplasm and through the outer membrane to the extracellular face by the Lpt protein complex, where lipid A becomes a component of the outer face of the outer membrane with the polysaccharide displayed on the surface of the cell.

The tendency for genes responsible for production of a bacterial polysaccharide to be organized as a single, continuous operon, especially in the case of O-antigens and CPS has greatly facilitated the transfer of polysaccharide coding loci from their native species into a heterologous host, typically *E. coli.* Early methods generally centered around the generation of a cosmid library from fragmented genomic DNA, followed by screening of individual cosmids at the genomic or phenotypic level to locate clones conferring production of the polysaccharide of interest. This approach has been used to produce a variety of O-antigens from Gram-negative organisms including *Pseudomonas aeruginosa, Salmonella typhimurium* and *Yersinia enterocolitica* in an *E. coli* strain background (Goldberg et al. 1992). A similar approach has also been employed for the production of CPS from the Gram-positive organism *Streptococcus pneumoniae* in the Gram-positive host *Lactococcus lactis* (Gilbert et al. 2000). Cloning of sequenced, annotated polysaccharide biosynthetic loci has enabled production in *E. coli* of polysaccharides from diverse Gram-negative species such as *Bukholderia pseudomallei* (Garcia-Quintanilla et al. 2014) and *Francisella tularensis* (Cuccui et al. 2013). A further advance was the recent demonstration that various CPS structures from the Gram-positive bacterium *S. pneumoniae* could be produced in a Gram-negative host, namely *E. coli*, using the *en bloc* transfer of the entire CPS coding locus (Kay et al. 2016; Price et al. 2016). The recombinant CPS structures are produced essentially as an O-antigen in *E. coli*, and some features of processing appear to be borrowed from the host, including the action of the O-antigen ligase WaaL in attaching the polymerized polysaccharide to the outer core on lipid A, and subsequent transport to the outer surface of the cell. These findings demonstrated an unexpected cross-compatibility between systems from two disparate sources, and highlighted the mechanistic similarity of CPS biosynthesis in Gram-positive bacteria and O-antigen biosynthesis in Gram-negative bacteria.

Recently, the wide availability of whole-genome sequences and a thorough understanding of the mechanisms behind bacterial polysaccharide biosynthesis has led to a more informed approach to the production of heterologous polysaccharides. A recent study produced two different *Staphylococcus aureus* CPS structures by expressing combinations of *P. aeruginosa* and *S. aureus* glycosyltransferases in *E. coli*, with sugar precursors provided by a combination of *P. aeruginosa* enzymes along with native enzymes in the *E. coli* host. The resulting glycans were confirmed by MALDI-TOF/TOF tandem mass spectrometry analysis as having the same structure as the native CPS, and were recognized by capsular serotype-specific typing antiserum (Wacker et al. 2014). Hence, bacterial glycosyltransferase enzymes may be regarded as modular entities defined only by function, opening up a new approach to polysaccharide bioengineering in host species such as *E. coli.* This insight also facilitates the engineering of bacterial glycans in cases where information regarding the biosynthesis of a target polysaccharide (and/or its intermediates) is incomplete or incompatible with further processing as a result of assembly on a lipid other than Und-PP. For example, the Vi polysaccharide of *Salmonella enterica* serovar Typhi is currently licensed as a purified polysaccharide vaccine for typhoid fever, but represents an interesting candidate for further development as a glycoconjugate. Unfortunately, recombinant production of this polysaccharide is challenging because the lipid on which it is assembled in the native host is not currently known. To circumvent this issue, Wetter et al. modified the *E. coli* O121 O-antigen, a structure that is well known to build on Und-PP, to resemble the Vi polysaccharide. Following transfer of the resulting Vi-like polysaccharide to a carrier protein, a glycoconjugate was produced that elicited antibodies that were immunoreactive with *E. coli* O121 LPS (Wetter et al. 2013).

## Bioengineering of eukaryotic polysaccharides on the LPS core in bacteria

The ability to expand the bacterial polysaccharide production system to engineer structures beyond prokaryotic polysaccharides is crucial if this approach is to become broadly applicable and useful. Several human-like glycans have been assembled on a truncated LPS outer core structure. Typical mutations involve the disruption of the second glycosyltransferase enzyme of the outer core, resulting in an intact lipid A molecule, coupled to a complete inner core structure, but with only a single glucose residue from the outer core added to the second heptose residue of the inner core (see Fig 2.). This exposed glucose then becomes the attachment site for recombinant glycans, while the Lpt export system translocates the resulting LOS structure to the surface of the cell, ensuring the recombinant glycan is exposed (Merritt et al. 2013).

The human glycosphingolipid globotriaosylceramide (Gb_3_) is the receptor for Shiga-toxin (Stx), a potent AB5 toxin produced by pathogenic species such as *Shigella dysenteriae* and *E. coli* O157. This receptor is composed of a trisaccharide, Gal(α1–4)Gal(β1–4)Glc, and is present on many eukaryotic cell types, but is found at the highest concentrations in renal tissue and in microvascular endothelial cells (Paton et al. 2000). An analogous structure to the Gb_3_ receptor is produced by *Neisseria spp.* as a component of LOS and is representative of a common strategy employed by mucosal pathogens whereby surface display of host glycan epitopes aids immune evasion (Cress et al. 2014). Expression of the glycosyltransferases LtxC from *Neisseria meningitidis*, and LtxE from *Neisseria gonhorroeae* in *E. coli* resulted in the production of a novel LPS-associated Gb_3_ polysaccharide structure. When administered to mice, the engineered *E. coli* were found to protect against challenge with a Shiga-toxin producing *E. coli* (STEC) strain, suggesting an effective molecular mimic of the toxin binding site had been recreated that sequestered the secreted toxin (Paton et al. 2000). An analogous approach has been used to engineer *E. coli* cells that express molecular mimics for other receptors implicated in bacterial toxin binding: globotetraosylceramide (Gb_4_), and the gangliosides GM_1_ and GM_2_ (Focareta et al. 2006; Hostetter et al. 2014). These engineered bacterial strains have also proven efficacious in animal models for the treatment of toxin-associated bacterial infections such as cholera and STEC.

A similar approach was used to produce the ganglioside GM_3_ epitope, NeuNAcα(2,3)Galβ(1,4), as an attachment to the exposed glucose residue of truncated lipid A (Ilg et al. 2010). This feat was accomplished by expressing the *Neisseria* enzymes SiaB, a CMP-sialic acid synthetase, together with the galactosyltransferase LgtE and the sialic acid transferase Lst, which together generated a GM_3_-like structure that was displayed on the surface of the cell. This strain may be useful for investigating the effects of sialic acid-containing bacterial LOS structures and their role in development of post-infection autoimmune diseases such as Guillain-Barre syndrome. Other human-like glycans with a role in bacterial attachment have also been expressed in *E. coli*, including fucosylated oligosaccharides: the blood group H, Lewis X (Le^X^) and Lewis Y (Le^Y^) antigens (Yavuz et al. 2011) and poly-N-acetyllactosamine (Mally et al. 2013). Fucose is a common component of human glycans, and is thought to play a role in the binding of various pathogenic bacteria including *P. aeruginosa* and *C. jejuni*, and it is envisioned that these strains may prove useful for studying specific bacterial interactions with human receptors, as well as informing the design of competitive inhibitors for novel probiotic-based therapies.

A further example of a eukaryotic glycan that may also be produced as a bacterial mimic is polysialic acid (PolySia), a linear homopolymer of α-2,8-linked sialic acid residues. In humans, this glycan is most notably found as an elaboration of the *N*-linked glycan on neural cell adhesion molecule (NCAM), but is also expressed by *E. coli* K1 and *N. meningitidis* group B as the K1 capsule and CPS A, respectively (Moe et al. 2009). Owing to its occurrence on these pathogens as well as its enhanced expression on some malignant tumors (Livingston et al. 1988; Komminoth et al. 1991), PolySia represents an intriguing target for vaccine or therapeutic antibody development. By expressing a combination of glycosyltransferases from *N. gonorrhoeae, C. jejuni* and *E. coli*, Valentine and co-workers were able to produce PolySia directly on the LPS core of an *E. coli* strain not normally capable of synthesizing this structure. Interestingly, where the aforementioned GM_3_ production study supplied NeuAc via the growth medium and relied on a single synthetase enzyme to convert the sugar into the nucleotide activated form CMP-NeuAc (Ilg et al. 2010), here the authors reconstituted the entire biosynthesis pathway capable of converting the readily available housekeeping sugar UDP-GlcNAc into CMP-NeuAc (Valentine et al. 2016), highlighting the flexibility and versatility of bacteria as hosts for glycoengineering.

## Bioengineering of eukaryotic polysaccharides on the lipid anchor Und-PP in bacteria

Because direct conjugation to the LPS core is not always possible or desirable, alternative sites for polysaccharide assembly have also been explored such as the common lipid anchor Und-PP. In *E. coli* K-12, the ECA and O-antigen biosynthesis pathways involve installation of a GlcNAc residue on Und-PP by an initiating glycosyltransferase called WecA. By introducing glycosyltransferases from the *Haemophilus influenzae* LOS biosynthesis pathway that were capable of modifiying this Und-PP-linked GlcNAc in the recombinant system, a tetrasaccharide resembling the Le^X^ antigen (minus the fucose residue) was assembled on Und-PP (Hug et al. 2011). The use of this lipid as a carrier enabled subsequent conjugation of the glycan to a protein using an oligosaccharyltransferase-mediated mechanism that is described in greater detail below. To complete the Le^X^ structure, the purified glycoconjugate was subjected to in vitro enzymatic elaboration to add the fucose residue (Hug et al. 2011). The use of engineered bacteria to produce Le^X^ containing glycoproteins is significant because these proteins are known to function as immunomodulatory molecules (Atochina et al. 2001; van Die et al. 2003; Srivastava et al. 2014), and have been shown to ameliorate symptoms associated with autoimmune disorders in animal models (Atochina and Harn 2006).

Another human-like glycan produced in a similar manner is the Thomsen-Friedenreich antigen (T antigen), a Galβ1-3GalNAc disaccharide. Valentine et al. used UndPP-linked GlcNAc as a primer for producing the T antigen disaccharide (Valentine et al. 2016). This was accomplished by addition of two heterologous glycosyltransferases and a nucleotide sugar epimerase to ensure availability of the required substrate UDP-GalNAc. Because T antigen is overexpressed on a number of malignancies including breast, colon, prostate and stomach cancer (Heimburg-Molinaro et al. 2011), recombinant biosynthesis could yield highly immunogenic glycoconjugates that elicit antibodies against this important glycan epitope.

A final example of engineering human-like glycans in a bacterial host involved the bottom-up creation of a eukaryotic *N*-glycan biosynthesis pathway. Specifically, the conserved core of all human *N*-glycans, the oligosaccharide Man_3_GlcNAc_2_, was successfully produced on Und-PP by co-expression of four eukaryotic glycosyltransferases, including the yeast uridine diphosphate-*N*-acetylglucosamine transferases Alg13 and Alg14 and the mannosyltransferases Alg1 and Alg2 (Valderrama-Rincon et al. 2012). By including a bacterial oligosaccharyltransferase PglB from *C. jejuni*, glycans were successfully transferred to eukaryotic target proteins as discussed below. The Man_3_GlcNAc_2_ structure has been shown to be the minimal structure required for efficacy of a glycoprotein therapeutic (Van Patten et al. 2007), and is the predominant glycoform conjugated to proteins expressed in a baculovirus host system. Furthermore, as the conserved core of human *N*-glycans, this structure has enormous potential as a precursor for further modification, either in vivo or in vitro.

### Glycoprotein expression in bacterial hosts: current applications and future opportunities

The above findings demonstrate the remarkable versatility of bacterial systems for the biosynthesis of a vast array of carbohydrate structures. However, to exploit the full potential of carbohydrates, it is often necessary to conjugate these structures to additional biomolecules such as proteins. Two different mechanisms are responsible for making the majority of proteins that become covalently modified with sugar molecules (*i.e.*, glycoproteins). These mechanisms are defined based on the amino acid residue onto which the glycan is installed. In *N*-linked glycosylation, the glycan is attached to the nitrogen atom of an asparagine residue, while in O-linked glycosylation the sugar moiety is attached to the oxygen atom of either a serine or a threonine side chain. While both types of glycosylation were long believed to occur exclusively in eukaryotes, multiple bacterial machineries for the generation of both types of modifications have been discovered over the last 15 years. These bacterial glycosylation systems, or hybrids thereof, have opened the door to using bacteria for the production of two important classes of glycoproteins: (1) glycoconjugate vaccines, whereby immunogenic carbohydrates from pathogens including bacteria and viruses are linked to proteins; and (2) therapeutic proteins that are glycosylated in their natural form and require the modification for full function, for example, monoclonal antibodies.

Glycoconjugates are amongst the most successful vaccines generated to date, eliciting a robust T-cell dependent immune response and conferring protection across all age groups (Vella and Pace 2015). For three important bacterial pathogens in particular, *H. influenza* type B (Hib), S. *pneumoniae* and *N. meningitidis*, glycoconjugates have proven to be highly effective in countries where they have been introduced (Ladhani 2012; Grijalva et al. 2007). The standard production method for these conjugates involves the separate generation and purification of the protein and the carbohydrate moiety, chemical activation thereof, and conjugation as well as subsequent purification of the resulting glycoprotein (Lees et al. 2008). Even though it is an established and accepted method, there are several drawbacks to this approach. Firstly, it requires culturing large volumes of a pathogenic species of interest for the generation of the native carbohydrate, followed by harvesting and purification of the carbohydrate. Depending on the biosafety level of the species of interest, as well as the ease of culturing, this step can present a major hurdle regarding the expansion of the technique to novel pathogenic species. Secondly, the activation and chemical conjugation steps required to couple the glycan to the carrier protein can be technically challenging and inefficient, resulting in low yields, as well as a heterogeneous population of glycoproteins with different numbers of target glycans attached at different locations throughout the protein. Therefore, alternative methods for generating glycoconjugates that overcome some of these limitations are desired.

In addition to glycoconjugate vaccines, many proteins of therapeutic interest are also glycoproteins. In fact, 70% of therapeutic proteins approved by regulatory agencies or currently in clinical and preclinical trials are decorated with glycans in their native form (Sethuraman and Stadheim 2006). Historically, this has limited the use of *E. coli* to proteins and peptides that are not natively glycosylated such as insulin and homologues thereof or to those that are natively glycosylated but are functional without the addition of the glycan moiety, such as human growth hormone (hGH) and interferon α (Ferrer-Miralles et al. 2009). It should be pointed out that these proteins often require additional post-translational modifications such as the addition of polyethylene glycol (PEG) to increase serum half-life (Bailon and Won 2009). While some notable breakthroughs have been made (Valderrama-Rincon et al. 2012), the routine use of *E. coli* as a production platform for therapeutic glycoproteins and glycopeptides requires further engineering of glycosylation pathways in this host.

### *N*-linked glycoprotein expression in bacteria

The discovery of an *N*-glycosylation machinery in the human intestinal bacterial pathogen *C. jejuni* (Szymanski et al. 1999) and the subsequent functional transfer of the complete machinery into the more tractable species *E. coli* (Wacker et al. 2002) demonstrated for the first time that bacteria could be an alternative source of recombinant *N*-glycoproteins. Subsequent studies showed that a single enzyme, an oligosaccharyltransferase named CjPglB (PglB from *C. jejuni)*, was responsible for transferring the glycan to the acceptor protein. Interestingly, this enzyme was shown to share sequence homology with the STT3 catalytic subunit of the eukaryotic oligosaccharyltransferase enzyme complex (Wacker et al. 2002). A functional study of the genes within the glycosylation locus demonstrated that the substrate glycan was assembled on the lipid carrier Und-PP (Linton et al. 2005), in a fashion similar to the O-antigen biosynthesis pathway present in many Gram-negative species of bacteria (Hug and Feldman 2011). It was further demonstrated that the CjPglB enzyme possesses remarkably relaxed glycan substrate specificity. That is, in addition to its native substrate oligosaccharide -- a heptasaccharide glycan with the structure diNAcBacGalNAc_5_GIc (Young et al. 2002) -- the enzyme was also able to recognize much larger polysaccharides such as structurally different bacterial O-antigens and transfer these to proteins (Feldman et al. 2005). Around the same time, a five amino acid glycosylation sequon for C*j*PglB was discovered (Kowarik et al. 2006), which could be engineered either into flexible secondary structures within a protein (Kowarik et al. 2006) or at either the N- or the C-terminus (Fisher et al. 2011). Altogether, these studies provided the requisite ingredients for making customized recombinant bacterial glycoproteins, where potentially any protein of interest could be modified with any glycan moiety at a desired position by co-expression of C*j*PglB, the glycan of interest assembled on Und-PP, and the desired acceptor protein modified to contain one or more glycosylation sequon(s) (Fig. 3).

**Figure 3.**
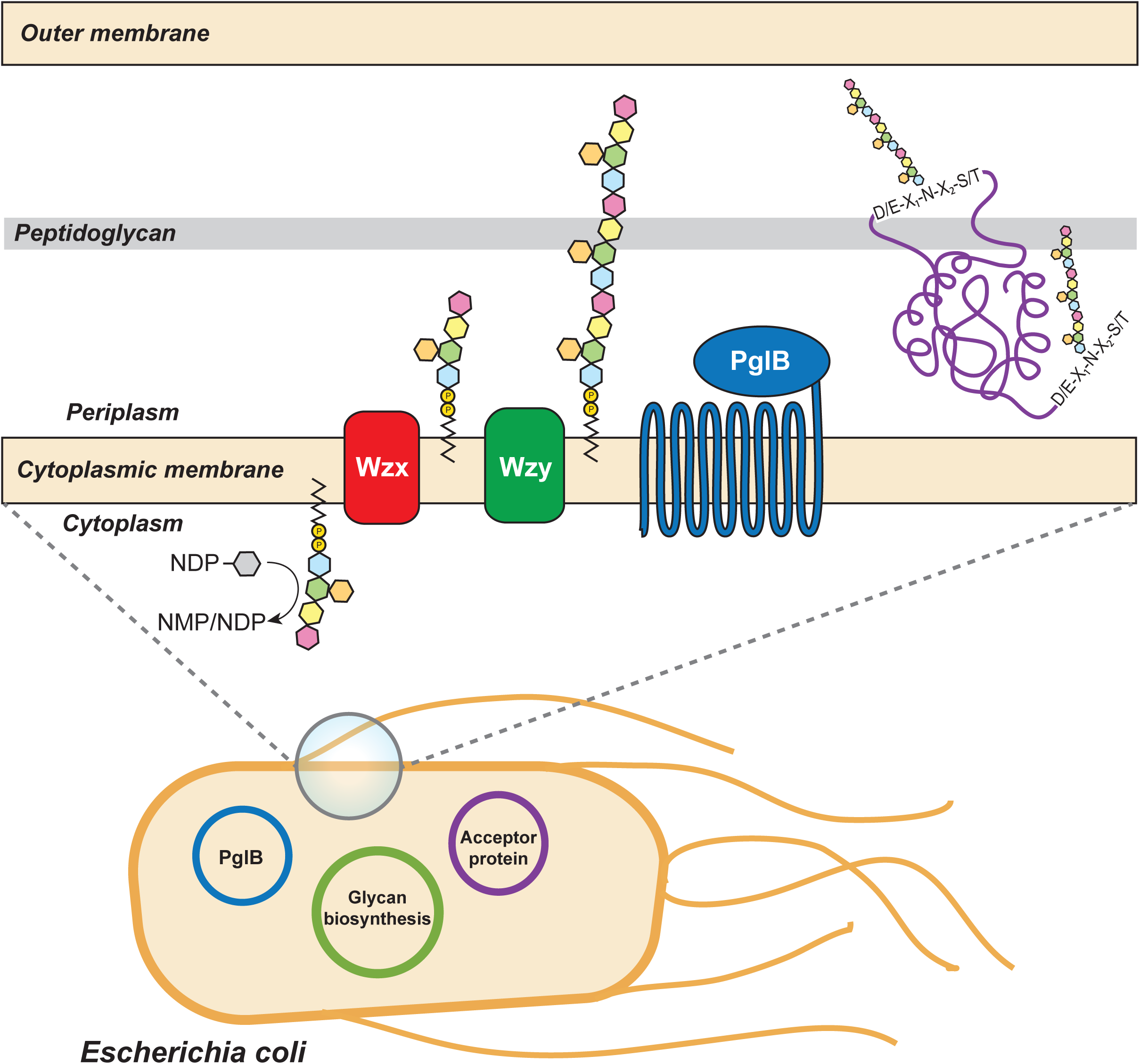
Recombinant protein glycosylation in *E. coli* using the bacterial oligosaccharyltransferase. Co-expression of three components is required for recombinant glycosylation in *E. coli:* (1) The glycan biosynthetic locus for the production of the carbohydrate of interest on the lipid carrier undecaprenol pyrophosphate; (2) the oligosaccharyltransferase (*e.g.*, C*j*PglB); and (3) the acceptor protein of interest that has been engineered with a signal peptide for export into the periplasm and an acceptor sequon (*e.g.*, D/E-X_1_-N-X_2_-S/T, where X can be any amino acid except proline) for glycosylation by the oligosaccharyltransferase. Sequons can be engineered into an exposed, flexible loop or at either the N- or the C-terminus of the protein. The glycoprotein can then be purified from the bacterial cells using standard methods.

## Customized *N*-glycoproteins produced recombinantly in *E. coli.*

To date, the predominant class of glycoproteins produced using the above components are conjugates in which bacterial surface glycan structures are site-specifically linked to immunogenic carrier proteins. In the majority of published cases, the glycans are O-antigen polysaccharides built on Und-PP (see above for in-depth discussion of different methods used for the recombinant production of these structures) and installed on the carrier protein by C*j*PglB. Table 2 summarizes the glycoconjugate vaccine candidates generated and tested to date. While multiple studies have demonstrated the generation of specific, and potentially protective antibody responses against *E. coli*-derived glycoconjugate vaccine candidates, it is particularly noteworthy that two have been successfully tested in Phase I trials. The first is a conjugate vaccine candidate against S. *dysenteriae* type 1 composed of the O-antigen glycan coupled to the exotoxin A of *P. aeruginosa* (EPA). Testing of this vaccine candidate in healthy adults at two different doses with or without co-adminstration of adjuvant revealed it to be well tolerated and capable of eliciting statistically significant antigen-specific humoral immune responses (Hatz et al. 2015). A second conjugate vaccine candidate comprised of the S. *flexneri* 2a O-antigen conjugated to EPA was also tested in healthy adults, with similar results regarding tolerance and immunogenicity (Riddle et al. 2016). Hence, recombinant production of glycogonjugates in *E. coli* appears to be a promising alternative to the traditional methods used for biomanufacturing conjugate vaccines.

**Table 2:**
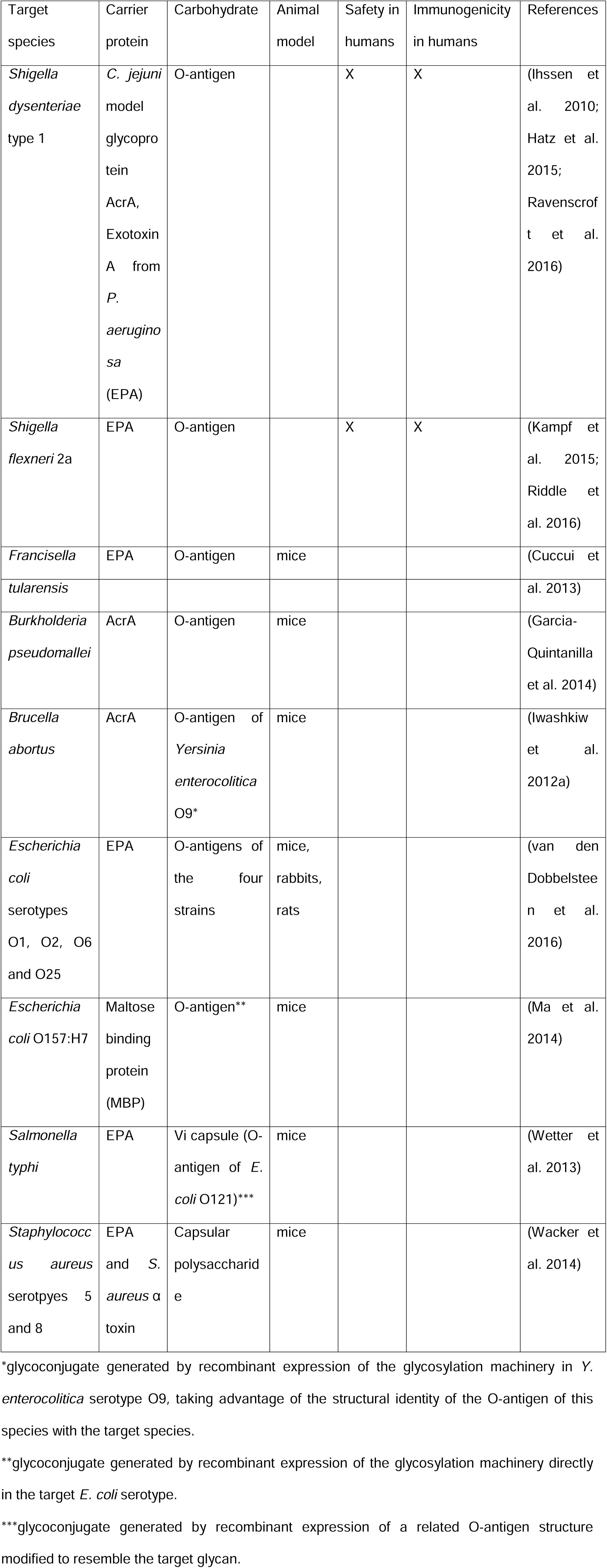
List of bacterial glycoconjugate vaccine candidates that have been produced using a bacterial glycoengineering approach.

Glycoconjugate proteins produced recombinantely in *E. coli* have found uses in other applications as well. For instance, bacterial glycoconjugates have been successfully used as diagnostic tools for human and bovine brucellosis (Ciocchini et al. 2013; Ciocchini et al. 2014) as well as for the Shiga-toxin-producing *E. coli* serotypes O157, O145, and O121 (Melli et al. 2015). Additionally, Shang and co-workers generated a glycoconjugate comprised of the maltose binding protein (MBP) and the *E. coli* O86:B7 O-antigen, which bears structural similarity to the blood group B antigen epitope. This glycoconjugate functioned as a ‘molecular sponge’ to lower the levels of blood group B antibodies in plasma without negatively affecting the clotting function of the plasma (Shang et al. 2016).

While there is a lot of promise for glycoconjugates where the sugar moiety is derived from an immunogenic bacterial glycan, these types of glycans will not be useful in applications where the goal is to install native, eukaryotic glycans onto therapeutic proteins. Several efforts have been made to leverage the bacterial protein glycosylation machinery for the generation of glycoproteins carrying mammalian glycans. Perhaps the most notable example is Valderrama-Rincon et al. who demonstrated the complete recombinant assembly and transfer to protein of the eukaryotic *N*-linked core glycan GlcNAc_2_Man_3_ (Valderrama-Rincon et al. 2012) (see above for description of the approach used for biosynthesis of the glycan). Transfer of the GlcNAc_2_Man_3_ glycan to asparagine residues in several different target proteins including the Fc domain of human immunoglobulin G (IgG) was achieved with C*j*PglB, which as mentioned above has fairly relaxed specificity towards the glycan substrate. One can envision an extension of this glycan, either *in vivo* or potentially *in vitro*, to generate additional structures found in mammalian N-glycans. In an alternative approach, post-processing of a purified pre-form of the glycoconjugate outside of the bacterial cell can be performed to generate the final product. For instance, the same GlcNAc_2_Man_3_ glycan structure was installed on a protein by a combination of recombinant *in vivo* glycosylation of the protein with the *C. lari* heptasaccharide glycan, GalNAc_5_GlcNAc, followed by *in vitro* enzymatic trimming of the glycan down to a single GlcNAc residue, and finally transglycosylation of the trimmed glycan with a preassembled Man_3_GlcNAc sugar to obtain the final structure (Schwarz et al. 2010). A similar combined method of in vitro and in vivo glycosylation and modification was used to install the blood group antigen Le^x^ on a protein (Hug et al. 2011). The recombinantely expressed tetrasaccharide GalNAc_2_Gal_2_ was produced on Und-PP in *E. coli* and this glycan was subsequently transferred *in vivo* to an acceptor protein using C*j*PglB. Following purification, *in vitro* fucosylation was performed to yield the final Le^x^ glycan on the protein. Although these combined *in vivo* and *in vitro* methods of glycoprotein biosynthesis are potentially less applicable to large scale production of glycoproteins, they nevertheless expand the range of glycan modifications on proteins, which may be beneficial for the generation of glycoproteins carrying sugars that are potentially too challenging for the expression and transfer *in vivo* alone.

## Expanding glycosylation through identification of alternative oligosaccharyltransferases

While C*j*PglB remains one of the best-characterized bacterial oligosaccharyltransferases, there are two main limitations that restrict its use for the coupling of designer glycans to acceptor proteins. First, compared to the canonical eukaryotic glycosylation sequon, N-X-S/T (where X can be any amino acid except proline), used by eukaryotic oligosaccharyltransferases, C*j*PglB requires an extended sequon (D/E-X_1_-N-X_2_-S/T) for the attachment of glycans to proteins (Kowarik et al. 2006). One consequence of this requirement is that, at a minimum, these five amino acids need to be engineered into the protein of interest either by addition of the residues as a terminal or internal tag or by changing of a native stretch of amino acids to render it a substrate for glycosylation. If these modifications are added to either of the termini, it can be speculated that this will not have a major impact on the overall structure and function of the protein. However, it may be desirable to engineer the site of glycan attachment into the protein, in which case these modifications may interfere with protein folding and/or function. Another consequence is that native *N*-glycoproteins of mammalian origin will need to have their shorter sequons extended to include a D or E in the −2 position to be glycosylated by C*j*PglB.

To address this limitation, several groups have used bioinformatics to identify orthologues of C*j*PglB, which were then functionally characterized in glyco-competent *E. coli* cells (Jervis et al. 2010; Ielmini and Feldman 2011; Schwarz et al. 2011b; Ollis et al. 2015; Mills et al. 2016). From these studies, oligosaccharyltransferases were identified from two species of *Desulfovibrio*, in particular, that did not require the negatively charged amino acid at position −2 and were therefore able to glycosylate the shorter eukaryotic N-X-S/T sequon (Ielmini and Feldman 2011; Ollis et al. 2015). Of these, only the PglB orthologue of *D. gigas* was able to modify the native QYNST sequon in the Fc domain of human IgG (Ollis et al. 2015), suggesting that additional factors govern acceptor-site specificity and must be satisfied to allow for the installation of glycans onto shorter eukaryotic sequons. Additionally, the orthologue from *D. desulfuricans* showed markedly lower efficiency in transferring the *E. coli* O7 O-antigen polysaccharide (Ielmini and Feldman 2011), suggesting this enzyme may not be as flexible as C*j*PglB regarding the glycan structure. As no other polysaccharides were tested as substrates for the *D. desulfuricans* PglB, it is unclear whether the low efficiency of transfer the O7 O-antigen is specific to this substrate or an inherent property of the enzyme. The ability of the orthologues from *D. vulgaris* and *D. gigas* to transfer mono- and polysaccharides was not tested, so it remains unclear whether these enzymes may be useful in the generation of custom glycoconjugates.

In parallel to the functional characterization of C*j*PglB orthologues, a directed evolution approach has applied to C*j*PglB with the goal of relaxing the acceptor-sequon specificity. Using the crystal structure of the closely related PglB enzyme of *C. lari* (Lizak et al. 2011) as guidance, combined with a high-throughput genetic screen using a secreted acceptor protein, a library of C*j*PglB mutants was screened for the ability of the enzyme to glycosylate non-canonical acceptor protein sites (Ollis et al. 2014). This screen identified three C*j*PglB variants that no longer required the negatively charged residue at the −2 position. The three mutants glycosylated a eukaryotic protein at its native N-X-S/T sequon, suggesting that these enyzmes may be useful for authentically glycosylating eukaryotic proteins and peptides. While the glycan specificity was not specifically tested, the fact that the mutants were derived from C*j*PglB suggests that the relaxed glycan specificity of the parent enzyme will remain.

A second limitation of C*j*PglB is the requirement of the native enzyme for an acetamido group at the monosaccharide that constitutes the reducing end of the oligo- or polysaccharide (Wacker et al. 2006). Many glycans of interest do not terminate in a glycan that conforms to this requirement, such as most capsular glycans of S. *pneumoniae* serotypes that terminate in either galactose or glucose residues (Bentley et al. 2006). While a natural variant among the orthologues of C*j*PglB enzymes from other species may lack this requirement, evidence for this has yet to be reported. In fact, two studies analyzing the protein *N*-glycan diversity within the *Campylobacter* genus and in one species of *Helicobacter* identified exclusively sugars containing an acetamido group at the reducing end (Jervis et al. 2012; Nothaft et al. 2012), suggesting that this is a shared feature among many of the bacterial species that possess protein *N*-glycosylation machineries. The same appears to be true for the sugar attached to an identified glycoprotein in *D. gigas*, which was *N*-glycosylated with a di-saccharide of GlcNAc and *N*-acetylallosamine (Santos-Silva et al. 2007). To address this issue, one study used structure-guided mutagenesis to engineer a C*j*PglB variant that was able to transfer two O-antigens from S. *typhimurium* which both contain non-acetylated sugars (galactose residues) at the reducing end (Ihssen et al. 2015). This work demonstrates that the glycan specificity of C*j*PglB can be engineered to a certain extent, and suggests that in the future it will be possible to transfer virtually any glycan to any protein using modified versions of C*j*PglB.

### Alternative routes for bacterial protein *N*-linked glycosylation

A novel family of bacterial enzymes has recently emerged that may be of potential use in bacterial glycoengineering. In contrast to the enzymes described in the previous section, these enzymes: (1) are active in the bacterial cytoplasm, not the periplasm; (2) use nucleotide-activated glycans instead of lipid-linked glycans as a substrate; and (3) recognize the shorter, bacterial N-X-S/T glycosylation sequon (McCann and St Geme 2014). The first member of the family was discovered in *H. influenzae*, and shown to be involved in the glycosylation of the high molecular weight adhesin protein HMW1 (Grass et al. 2003). The glycans attached to the adhesin protein were identified predominantely as hexose sugars, and glycosylation of the adhesin protein was demonstrated to be important for correct secretion of the adhesin as well as adhesion of the bacteria to airway epithelial cells (Grass et al. 2010). Further members of the family have been identified in several other species of bacteria (McCann and St Geme 2014), and in vitro experiments confirmed activity of the orthologues from *Y. enterocolitica* and *Actinobacillus pleuropneumoniae* (Schwarz et al. 2011a). The preferred substrate for the *A. pleuropneumoniae* enzyme (termed ApNGT) was demonstrated to be UDP-Glc (Schwarz et al. 2011a), and a downstream gene was shown to encode a glycosyltranferase enzyme that was able to extend the Glc moiety installed by ApNGT with further Glc residues. Additionally, when expressed in *E. coli*, ApNGT was shown to glycosylate recombinantly co-expressed auto-transporter proteins from the same species (the enzyme’s native substrate), as well as co-expressed human erythropoietin (EPO) and several native *E. coli* proteins (Naegeli et al. 2014). A polypeptide modified with a glucose moiety by ApNGT was also successfully elaborated through *in vitro* transglycosylation mediated by endoglycosidase enzymes (Lomino et al. 2013). This suggests that ApNGT and other enzymes from this family may be useful tools for installation of a priming glucose residue on proteins of interest followed by either *in vitro* or *in vivo* elaboration of the glycan. It can also be envisioned that directed evolution of the enzyme from this family may allow for the modulation of the carbohydrate specificity in a similar way to C*j*PglB.

### Customized *O*-glycoproteins produced recombinantly in *E. coli.*

In addition to the bacterial *N*-glycosylation mechanisms discussed above, pathways that lead to the modification of serine or threonine residues (*O*-linked glycosylation) have also been identified in several bacterial species. These mechanisms are more commonly found in bacteria than their *N*-glycosylation counterparts (Iwashkiw et al. 2013), and are currently being pursued for recombinant protein glycosylation. The following section will highlight similarities and differences between the *N*- and O-linked pathways.

Over the last decade, O-glycosylation machineries that share mechanistic similarities with the *N*-glycosylation pathways described above have been identified and characterized in several bacterial species (Iwashkiw et al. 2013). It was initially observed that the type IV pilus subunit protein PilA in *P. aeruginosa* strain 1244 was modified with a glycan in a manner dependent on the product of the gene adjacent to *pilA* named PilO/TfpO (Castric 1995). A similar machinery was identified in *N. mengingitidis*, whereby deletion of a gene termed *pglL* led to the loss of a carbohydrate moiety from the pilus subunit protein PilE (Power et al. 2006). Interestingly, both the *P. aeruginosa* PilO/TfpO and the *N. meningitidis* PglL proteins showed homology to O-antigen ligase proteins that are involved in transfer of the O-antigen subunit from the lipid carrier Und-PP onto the lipid A moiety during LPS biogenesis (Whitfield et al. 1997).

This suggested that these enzymes may use Und-PP-linked glycans as substrate. Analysis of the glycan structure present on *P. aeruginosa* PilA showed the presence of a single O-antigen repeat unit, further strengthening the hypothesis that Und-PP-linked glycans may be the substrate for this enzyme family (Castric et al. 2001). When PilO/TfpO and PilA from *P. aeruginosa* (or PglL and PilE from *N. meningitidis)* were recombinantly co-expressed in *E. coli* along with a Und-PP-linked oligo- or polysaccharide, transfer of the glycan to the pilin protein was observed (Faridmoayer et al. 2007). These results not only demonstrated recombinant activity of this new family of bacterial O-oligosaccharyltransferase enzymes, but also confirmed the substrate identity as Und-PP-linked glycans. Further analysis of the glycan specificity of PglL demonstrated a remarkable promiscuity with regards to the glycan. Diverse glycan structures were shown to be transferred to PilE by PglL in vivo including structures containing a Gal residue at the reducing end such as the S. *typhimurium* LT2 O-antigen and the di-saccharide-pentapeptide peptidoglycan building block, none of which are substrates for the *C. jejuni* oligosaccharyltransferase CjPglB (Faridmoayer et al. 2008). Additionally, in vitro glycosylation experiments revealed that the enzyme displayed flexibility toward the lipid carrier (Faridmoayer et al. 2008; Musumeci et al. 2013). Altogether, these characteristics suggest that this enzyme is a very promising tool for the generation of designer glycoproteins with *O*-linked sugars.

To date, however, the biotechnological use of this enzyme family has been hampered by one major bottleneck. Unlike in the case of CjPglB, there is a lack of a consensus sequon for glycosylation that would allow for the ‘tagging’ of any protein as a substrate for *O*-glycosylation. Analysis of the *O*-glycome of several organisms that possess PglL-like *O*-glycosylation systems identified multiple glycosylated proteins, and while these helped to determine that the amino acid residues around the glycan attachment site were rich in serine, proline and alanine, they did not reveal the presence of any consensus sequence (Vik et al. 2009; Iwashkiw et al. 2012b; Lithgow et al. 2014; Elhenawy et al. 2016). Towards a more universal glycosylation strategy, Qutyan and co-workers showed that a C-terminal fusion of *E. coli* alkaline phosphatase with the final 15 amino acids from the C-terminus of PilA was glycosylated by PilO/TfpO when expressed in *P. aeruginosa;* however, the observed glycosylation was not very efficient (Qutyan et al. 2010). Additionally, while it has been shown that PilO/TfpO has relatively relaxed specificity and was able to transfer multiple different serotype *O*-glycans of *P. aeruginosa* (DiGiandomenico et al. 2002), the enzyme was only able to transfer a single *O*-antigen subunit both in the native organism as well as recombinantly in *E. coli* (DiGiandomenico et al. 2002; Faridmoayer et al. 2007). Hence, alternative PilO/TfpO O-oligosaccharyltransferases will need to be identified or engineered for transferring longer polysaccharides, which are often desirable for glycoengineering purposes. This issue appears to have been solved recently by Pan and coworkers who reported the development and optimization of an O-linked ‘glycosylation tag’ consisting of an 8 amino acid motif flanked by two approximately 10 amino acid sequences containing mainly hydrophilic residues (Pan et al. 2016). This tag was successfully fused to both the N- and C-termini of three potential vaccine carrier proteins -- the cholera toxin B subunit, exotoxin A from *P. aeruginosa*, and the detoxified variant of diphtheria toxin CRM197 -- and glycosylated with two different sugars including the S. *typhimurium* LT2 O-antigen, which as discussed above is not a substrate for C*j*PglB. Recombinant O-glycoproteins produced with this method were tested in a series of animal experiments and elicited a glycan-specific antibody response (Pan et al. 2016). The ability to tag proteins for PglL-dependent O-glycosylation opens up this enzyme family for biotechnological applications, in particular in cases where the glycan of interest may not be an optimal substrate for *N*-glycosylation by C*j*PglB.

## Alternative routes for bacterial protein *O*-linked glycosylation

Many bacterial species possess *O*-glycosylated flagellar proteins, with the glycosylation patterns ranging from a single glycan at a single site to multiple glycans attached to different sites on the protein (Nothaft and Szymanski 2010). These glycans are installed in a processive manner, with individual glycosyltransferases adding the glycans sequentially to the protein. This mechanism is similar to the installation of *O*-linked glycans in eukaryotic mucin-like glycosylation (Kudelka et al. 2015). It could therefore be hypothesized that enzymes from these machineries could potentially be used/engineered to install mucin-like glycans on human proteins. The successful recombinant installation of the first monosaccharide of the core of human mucin-like glycan, a GalNAc residue, has been demonstrated in the cytoplasm of *E. coli* using a recombinantly expressed human GalNAc transferase enzyme (Henderson et al. 2011). However, no further elaboration of this priming glycan with other sugars has been demonstrated.

### Alternative therapeutic bacterial conjugates

Although some unconjugated polysaccharides are currently licensed as vaccines, they often elicit a T-cell independent immune response stimulated by the extensive cross-linking of receptors on the surface of B cells. As such, they are poorly immunogenic in children under two years of age and elderly patients, greatly limiting their usefulness (De Gregorio and Rappuoli 2014). While protein conjugation is the most widely studied approach to counter this problem, the field of bacterial glycobiology is opening up alternative approaches to boost the immunogenicity of carbohydrate epitopes.

One such approach is based on bacterial outer membrane vesicles (OMVs), which are small (20-200 nm in size) liposomes released from the outer membrane of nearly all Gramnegative bacterial species. These vesicles are non-replicating versions of their bacterial ‘parent’, and contain many of the same components as the bacterial outer membrane including membrane proteins, CPS, and LOS and LPS, as well as some of the luminal components of the bacterial periplasm (Kulp and Kuehn 2010). OMVs have garnered interest as vaccine candidates because vesicles from several bacterial pathogens have been shown to possess potent immunogenic capacities (Schild et al. 2008; Ellis et al. 2010; Alaniz et al. 2007). Intriguingly, OMVs appear to also possess intrinsic adjuvant properties, potentially removing the need to include adjuvants in the formulation (Sanders and Feavers 2011; Chen et al. 2010) OMVs derived directly from pathogenic *N. meningitidis* have been successfully incorporated into a commercial vaccine formulation, the recently licensed Bexsero (Holst et al. 2009; Gorringe and Pajon 2012). Native OMVs have been further engineered to carry additional immunogenic proteins, which are recombinantly displayed on the surface of the OMV through genetic fusion to outer membrane proteins or in the OMV lumen through periplasmic expression (Chen et al. 2010; Muralinath et al. 2011). Importantly, robust immune responses against these recombinant immunogens have been demonstrated (Chen et al. 2010; Muralinath et al. 2011). Three recent reports highlight a novel bacterial glycoengineering approach to OMV-based vaccines whereby immunogenic glycans are recombinantly displayed on the exterior of OMVs. The approach takes advantage of the following: (1) the fact that standard laboratory strains of *E. coli* have lost the ability to produce a native O-antigen glycan due to the insertion of an IS element in the second glycosyltransferase gene *wbbL* (Liu and Reeves 1994) while the rest of the mechanism including the flippase and ligase genes remain intact; (2) the ability to recombinantly express non-native polysaccharides in *E. coli;* (3) the fact that the O-antigen ligase WaaL has relative relaxed glycan specificity and will efficiently transfer engineered glycans from Und-PP to the lipid A-core in cells that lack the native O-antigen (Han et al. 2012); and importantly, (4) the recombinant O-antigen is efficiently transported to the cell surface and packaged into released OMVs. Using this approach, *E. coli-derived* glycosylated OMVs (glycOMVs) have been decorated with the O-antigens of eight Gram-negative bacterial species including *F. tularensis* (Chen et al. 2016), PolySia (Valentine et al. 2016), the CPS of S. *pneumoniae* serotype 14, and the N-linked heptasaccharide of *C. jejuni* (Price et al. 2016). Following immunization, the glycOMVs carrying the *F. tularensis* O-antigen were shown to elicit significant serum titers of class-switched, glycan-specific IgG antibodies in mice, and prolonged survival upon challenge with the highly virulent *F. tularensis* subsp. *tularensis* (type A) strain Shu S4 (Chen et al. 2016). Likewise, glycOMVs decorated with PolySia also elicited glycan-specific IgG antibodies in mouse immunization studies, and the serum antibodies had potent bactericidal activity, killing *N. meningitidis* serogroup B bacteria that possess a PolySia capsular glycan (Valentine et al. 2016). GlycOMVs carrying the S. *pneumoniae* serotype 14 CPS also elicited glycan-specific antibodies in mice, and the serum antibodies were shown to possess potent bactericidal properties when tested in an opsonophagocytic assay. In fact, the bacterial killing of the serum from mice vaccinated with the glycOMVs carrying the capsular glycan was as efficient as the serum from mice that had been vaccinated with the commercial glycoconjugate vaccine Prevnar13^®^ (Price et al. 2016). And finally, glycOMVs displaying the *C. jejuni* N-linked glycan were shown to lower levels *of C. jejuni* colonization in chickens by almost 4 logs (Price et al. 2016). The expansion of the technology to cover further species or serotypes is envisioned to be relatively straightforward, simply requiring the recombinant expression of a pathogen-specific glycan structure on the surface of *E. coli* cells.

A related approach to glycOMV vaccines is the development of whole-cell vaccines displaying recombinant glycan epitopes. This strategy also leverages the fact that recombinant polysaccharides assembled on Und-PP are often efficiently transferred to lipid A and displayed as recombinant chimeric LPS on the surface of Gram-negative bacteria. This approach has been evaluated using several different species of Gram-negative bacteria as hosts (S. *enterica* serovar Typhi, S. *enterica* serovar Typhimurium and *E. coli)* carrying biosynthesis gene clusters for immunogenic carbohydrates of S. *dysenteriae* serotype O1 (Xu et al. 2007), shiga-toxin producing *E. coli* serotype O111 (Wang et al. 1999), and *C. jejuni* (Nothaft et al. 2016). In contrast to glycOMV vaccine candidates, these whole-cell vaccine candidates replicate. While it is desirable to control their ability to replicate, a balance needs to be found between controlling the replication of the bacteria and ensuring they persist long enough in the vaccinated organisms to generate a desired immune response. Genetic inactivation of the *aroA* gene encoding a 5-enolpyruvylshikimate-3-phosphate synthetase, involved in the shikimate pathway that directly connects glycolysis to the synthesis of aromatic amino acids (Bentley 1990), is a commonly used strategy to attenuate live bacterial vaccine candidates. This is particularly useful in species of *Salmonella* as these mutants are able to grow in rich media *in vitro* but become self-limiting *in vivo*, where aromatic amino acids are not freely available (Ruby et al. 2012). However, recent data suggests that deletion of *aroA*, at least in S. *enterica* serovar Typhimurium, can lead to additional effects in cellular physiology that may have an influence on the behavior of the recombinant bacteria within the host (Felgner et al. 2016). Nonetheless, attenuated, glycan epitope-expressing bacteria offer an additional opportunity for glycoengineering of vaccine candidates, in particular in areas where minimal cost of production may be a priority, such as in poultry and other livestock vaccines.

### Concluding remarks and outlook

In summary, bacterial expression systems have been successfully used for the production of a variety of carbohydrate structures ranging from small secreted oligosaccharides to repeating polymers of high molecular weight, and spanning structures found in all kingdoms of life. Furthermore, the characterization of both *N*- and *O*-linked protein glycosylation systems in a variety of bacterial species has greatly enhanced the potential of bacterial systems for the generation of therapeutically relevant glycoconjugates. These bacterial conjugation systems have been employed to generate well-defined therapeutic compounds including the first conjugate vaccines produced entirely in bacteria as well as novel immunogenic entities such as glycosylated outer membrane vesicles. Two of these bacterially-derived glycoconjugates have recently undergone successful Phase I clinical trials, and new candidates are also emerging.

Owing to their versatility and ease of manipulation, bacteria are an ideal host for the production of a diverse array of structurally defined polysaccharides and glycoconjugates that will be of interest as medical and industrial products. Furthermore, the low costs associated with the culturing of bacterial strains, especially *E. coli*, opens up this technology to a far wider range of laboratories than existing chemical/chemoenzymatic synthesis methods or mammalian cell culture approaches. The findings from a recent report commissioned by the National Academy of Sciences states that ‘glycans play roles in almost every biological process and are involved in every major disease’ and further asserts that ‘the development of transformative methods for the facile synthesis of carbohydrates and glycoconjugates should be a high priority’ (National-Research-Council 2012). Bacterial glycoengineering represents an emerging field with the potential to play a major role in meeting these goals.

